# Effects of sheared chromatin length on ChIP-seq quality and sensitivity

**DOI:** 10.1101/2020.09.30.320697

**Authors:** Cheryl A. Keller, Elisabeth F. Heuston, Belinda Giardine, Maria R. Long, Amber Miller, Alexander Q. Wixom, Stacie M. Anderson, David M. Bodine, Ross C. Hardison

## Abstract

Chromatin immunoprecipitation followed by massively parallel, high throughput sequencing (ChIP-seq) is the method of choice for identifying, on a genome-wide scale, the segments of DNA bound by specific transcription factors (TFs) or in chromatin with particular histone modifications. However, the quality of ChIP-seq datasets vary over a wide range, with a substantial fraction being of intermediate to poor quality. Such experimental variability can lead to many false positives or false negatives, impairing the ability to interpret the data. Thus, it is important to discern and control the factors that contribute to variation in ChIP-seq. In this study, we focus on the sonication step to produce sheared chromatin, a variable controllable by the user and applicable to all ChIP-seq protocols. We systematically varied the amount of shearing of fixed chromatin from a mouse erythroid cell line, carefully measured the distribution of resultant fragment lengths using the Agilent Bioanalzyer 2100, and then immunoprecipitated these batches of chromatin using highly specific antibodies against either TAL1 or CTCF. We found that the level of sonication, which was affected by both the number of sonication cycles, as well as the starting cell number, had a pronounced impact on the quality of resulting ChIP-seq signals. Specifically, over-sonication led to degradation of quality (e.g. increased background and reduction in signal), while the impact of under-sonication of chromatin differed between the two transcription factors, leading to the loss of sites occupied by TAL1 but not those bound by CTCF. We leveraged these findings to produce a set of CTCF ChIP-seq datasets in primary hematopoietic progenitor cells, including several rare cell types. Together, these results suggest that the amount of sonication is a key variable in success of ChIP-seq experiments, and that carefully monitoring the level of chromatin sonication is one way to improve ChIP-seq quality and reproducibility, which in turn facilitates low input ChIP-seq in rare cell types.

## Introduction

Chromatin immunoprecipitation followed by massively parallel, high throughput sequencing (ChIP-seq) has been used extensively to produce thousands of genome-wide maps of DNA segments bound by specific transcription factors (TFs) or in chromatin with particular histone modifications. However, these ChIP-seq datasets vary widely in quality. A uniform analysis of vertebrate transcription factor ChIP-seq datasets in the Gene Expression Omnibus (GEO) repository found that a substantial fraction were of intermediate to poor quality, and many of the control datasets (e.g. IgG and mock immunoprecipitations) displayed enrichments similar to the experimental datasets, indicating that many ChIP-seq results are not substantially different from the negative controls (Marinov et al. 2014). An independent assessment of reproducibility in ENCODE ChIP-seq datasets found that while almost half the datasets had good agreement in peak calls across replicates, almost one-third (18/57) were dissimilar between replicates (Devailly et al. 2015). The variation in quality of ChIP-seq datasets is well-known, and efforts have been made to communicate the quality to potential users. A set of quality metrics were defined (Landt et al. 2012), and these are used to evaluate datasets within ENCODE and frequently in individual labs. Application of these and additional considerations has led to the introduction of audit “flags” at the ENCODE data portal (Davis et al. 2018). For the 7547 ChIP-seq datasets across four species (as of July 14, 2020), about 300 red flags and about 3000 orange flags were given (multiple flags can be assigned to one dataset), illustrating a serious, but not crippling, issue. These studies of reproducibility and data quality all illustrate the variable quality in ChIP-seq datasets. Such experimental variability can lead to increased costs due to failed experiments, and it can lead to misinterpretation of data when the failures or very low quality of datasets are not recognized. Thus, it is important to discern and control the factors that contribute to variation in ChIP-seq.

Some of the factors that affect quality and reproducibility of ChIP-seq data are largely outside the control of the experimentalist. The major limiting factor in conventional ChIP-seq is the availability of highly specific antibodies. While many manufacturers make claims of “ChIP-seq validated” antibodies, many antibodies do not produce high quality ChIP-seq data, resulting in a high failure rate using commercially available antibodies against a diverse spectrum of TFs (Savic et al. 2015).

Additional variables that influence the success of a ChIP-seq experiment include the abundance of the target proteins and their accessibility in chromatin to antibodies. Assays of modified histones in chromatin are often highly sensitive and reproducible, likely due to the availability of good quality antibodies as well as their high expression levels and tight association with chromatin. By comparison, the level of expression of sequence-specific TFs varies among factors as well as across cell types. Further, the extent of interaction with chromatin varies depending on its binding site and whether or not it binds DNA directly or via DNA-associated factors.

Variables such as antibody quality or antigen abundance and accessibility can sometimes be controlled, but only by dedicated effort focused on the TF or epigenetic feature of interest. Other variables that are part of the preparative procedure, such as specific chromatin fixation and sonication conditions, are controlled by the experimentalist, and they can be optimized to improve sensitivity and reproducibility of chromatin immunoprecipitation (Khoja et al. 2019). Previous work has indicated a recommended size range between 100 and 300 bp (Landt et al. 2012; Browne et al. 2014), while other work suggests a wider size range of 100 to 600 bp (Diagenode 2012). Thus, the optimal size for ChIP-seq currently is not clear, and a convenient means for achieving an optimal size range has not been described.

In this study, we systematically evaluate how the extent of chromatin sonication affects ChIP-seq quality and success rate for two different sequence-specific TFs, CTCF and TAL1, in mouse erythroid cells. We further develop a case that the distribution of sizes of chromatin fragments after sonication is a critical, controllable aspect of ChIP-seq experiments. Finally, we leveraged these findings to produce a set of CTCF ChIP-seq datasets in primary hematopoietic progenitor cells, including several rare cell types such as hematopoietic stem and progenitor cells, and lineage-restricted progenitor populations.

## Results

To evaluate systematically how chromatin sonication affects ChIP-seq quality and success rate, we used the Diagenode Biorupter 300 to shear fixed chromatin from batches of 50M and 20M mouse erythroid G1E-ER4+E2 cells to varying degrees **(Supplemental Figure S1)**, assayed the extent of shearing using the Agilent Bioanalzyer 2100, and then subjected the chromatin to immunoprecipitation and sequencing using antibodies against either CTCF or TAL1 (**Figure 1A-D, Supplemental Table S1**), for which binding sites are well known.

**Figure 1:**
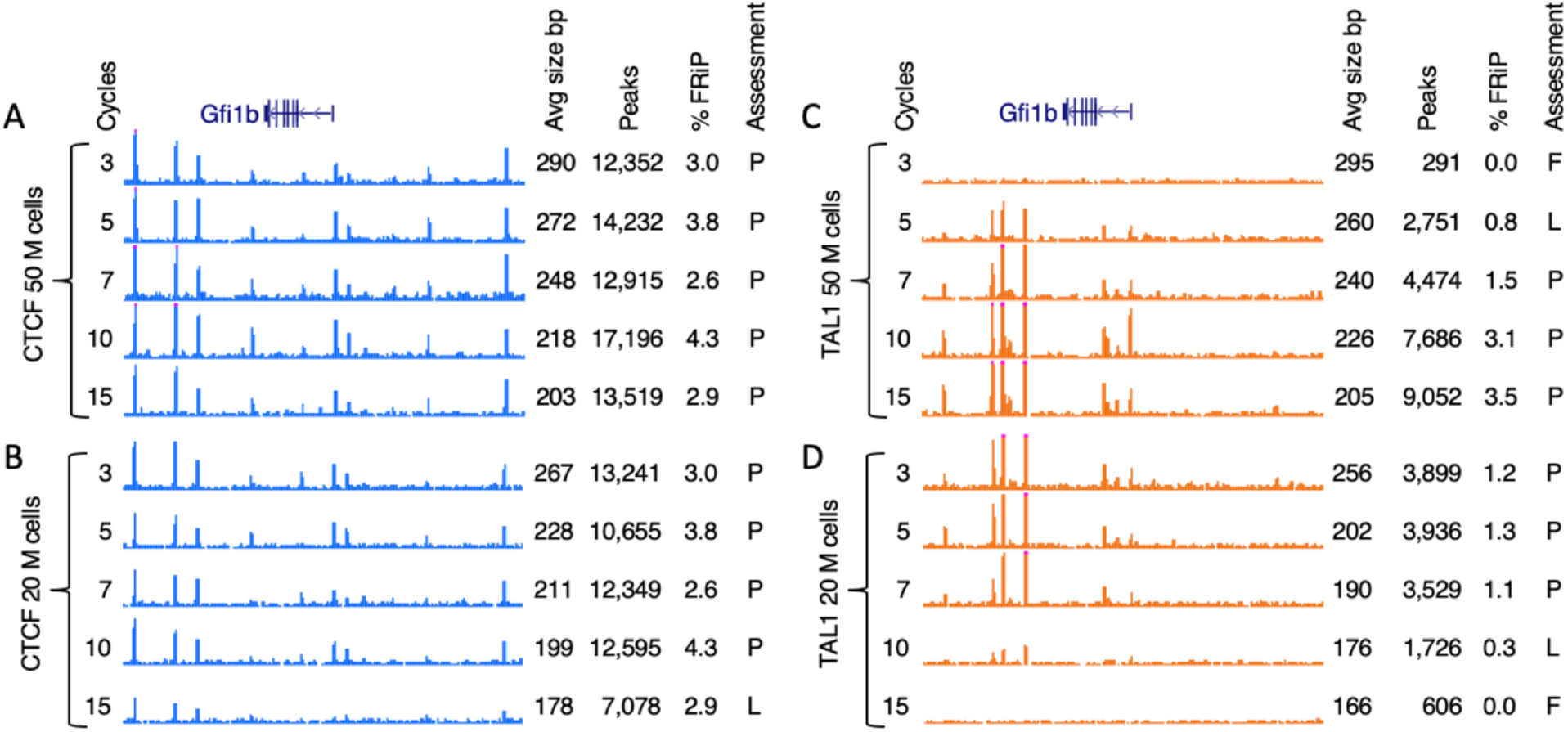
ChIP-seq signal tracks at the *Gfi1b* locus for **A**. CTCF 50M cells, **B**. CTCF 20M cells, **C**. TAL1 50M cells, and **D**. TAL1 20M cells. P = pass, L = low pass, F = fail.

The resulting ChIP-seq patterns revealed a striking dependence on numbers of sonication cycles. While many CTCF ChIP-seq samples (**Figure 1A, B**) showed the expected peaks at the illustrative locus *Gfi1b*, the sample from 20M cells sonicated for 15 cycles showed a low signal-to-noise (**Figure 1B**). Good quality TAL1 ChIP-seq data were obtained for most of the samples, but the pattern of failures was more complex, with poor results obtained at low cycle numbers for 50 M cells and at higher cycle numbers for 20M cells (**Figure 1C, D**).

We hypothesized that the lower quality datasets may result from the impact of the different cell numbers and sonication conditions on the resulting sizes of the chromatin. To interrogate the relationship between chromatin length and ChIP-seq success, we used the Agilent 2100 Bioanalyzer results to determine the average size of unenriched chromatin between 100-500 bp for each sample (**Figure 1, Supplemental Table S1, Supplemental Figure S2)**. This range was selected in order to standardize the measurement scheme, because large molecular weight heterochromatin could skew the average when chromatin size is measured over the entire range. Further, fragments outside of this size range are less likely to be sequenced when using the Illumina platform for sequencing the DNA after chromatin immunoprecipitation. We stress that the average chromatin fragment sizes measured in this manner are not the same as the average library size of the completed library or the library insert sizes deduced from the patterns of mapped reads after sequencing. The library sizes reflect the distribution of DNA fragments that were selected during the library preparation protocol; they are not the average size of the pool of unenriched chromatin. In other words, the DNA fragments measured by sequencing library size are the subset of immunoprecipitated chromatin that are most favorable for sequencing, whereas the measurements used in our study are those of the chromatin input for immunoprecipitation, which are controlled by the experimentalist.

We tested the hypothesized impact of chromatin size on success rate of ChIP-seq in several ways. First, we categorized the success or failure of each ChIP-seq experiment by inspection of the signal tracks in loci for which the CTCF and TAL1 patterns have been examined in prior work. For instance, the *Gfi1b* locus (**Figure 1A-D**) has been studied extensively by ChIP-seq and genetic experiments (Wilson et al. 2010; Dogan et al. 2015; Wilson et al. 2016; Xiang et al. 2020). ChIP-seq patterns with good signal-to-noise ratios and concordance with prior knowledge were classified as “Pass”, those with some peaks present but missing others were classified as “Low pass”, and those with almost no peaks were classified as “Fail” (**Figure 1A-D**). These subjective inspections were consistent with the objectively determined numbers of peaks called by MACS in each sample, with the experiments assessed as “Pass” having more peaks. (**Figure 1A-D, Materials and Methods**).

We found that, as expected, the chromatin size is inversely proportional to the numbers of cycles of sonication that were used for generating both the CTCF ChIP-seq datasets (R= -0.97 for 50M cells, R= -0.93 for 20M, **Figure 2A**) and TAL1 ChIP-seq datasets (R= -0.94 for 50M cells, R= -0.85 for 20M, **Figure 2B**). The number of cycles required to reach a given average chromatin size depended on the number of cells in the starting sample, with consistently more cycles of sonication needed to break chromatin to a given size when more cells are being processed. One would also expect sonication behavior to vary with different cell types and fixation methods, so in practice, the number of cycles to obtain a particular range of chromatin needs to be determined empirically. A procedure for determining the optimal number of cycles to obtain a good fragment size range is given in **Supplemental Figure S1**.

**Figure 2.**
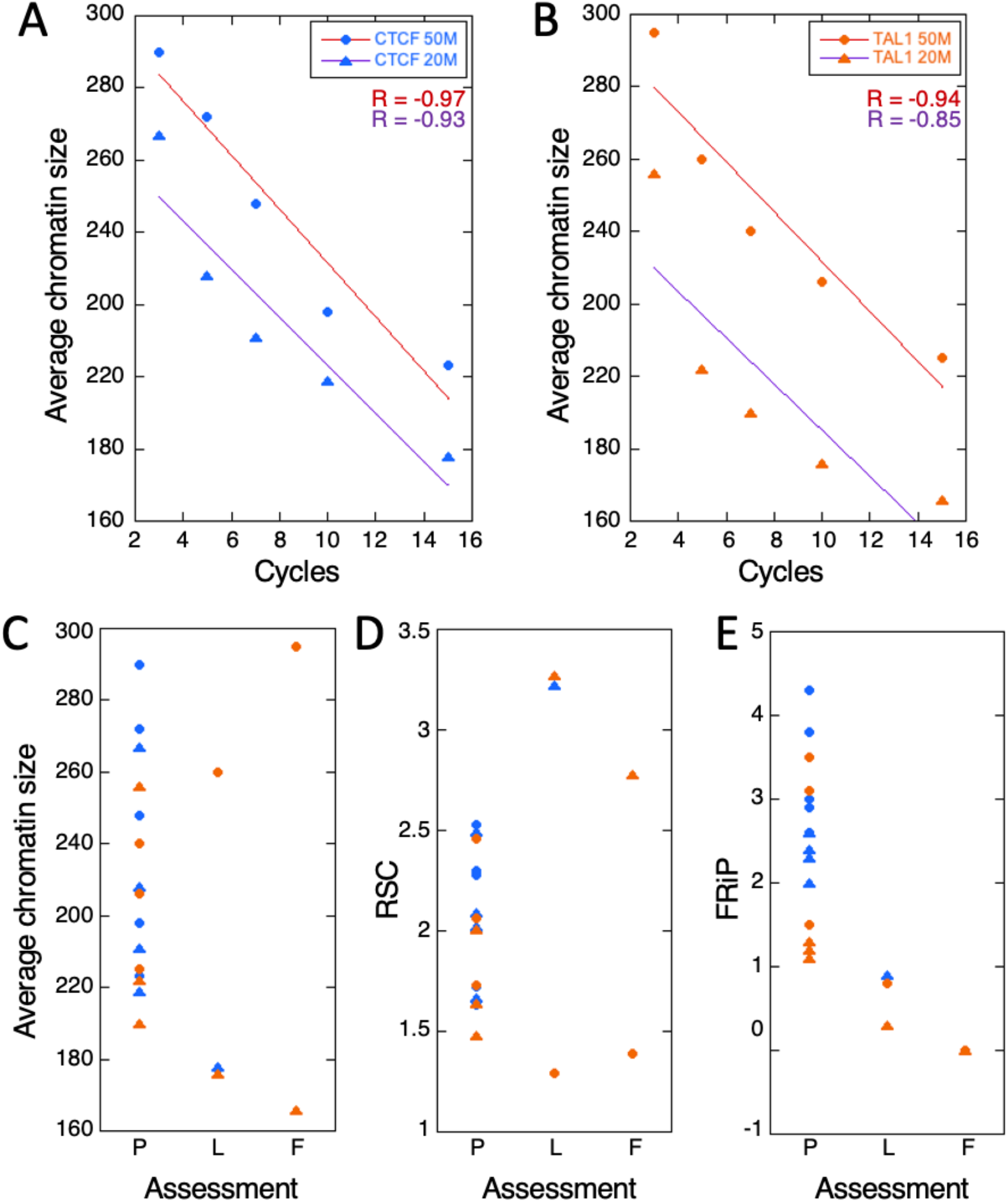
Effect of sonication cycles on chromatin length, and assessment of ChIP-seq success as related to objective quality metrics. **A**. Average chromatin length vs cycles, CTCF, **B**. Average chromatin length vs cycles, TAL1, **C**. Average chromatin length classified by subjective assessment, **D**. RSC classified by subjective assessment, **E**. FRiP scores (as percentage) classified by subjective assessment. CTCF = blue, TAL1 = orange, 50M = circles, 20M = triangles, P = pass, L = low pass, F = fail.

We then examined the relationship between the subjective success calls and the input chromatin size, and found that the successful ChIP-seq experiments all came from sheared chromatin whose size was within a fairly defined range (about 190 to 290 bp; **Figure 2C**). Samples for which the ChIP-seq failed or marginally passed tended to have chromatin sizes outside this range. These results support our hypothesis and indicate that chromatin size may be related to success frequency of ChIP-seq.

Next, we turned to objective quality metrics of each ChIP-seq experiment, specifically the Relative Strand Correlation (RSC) values and the Fraction of Reads in Peaks (FRiP) scores (Landt et al. 2012). Again, the samples with ChIP-seq patterns that passed our subjective visual inspection tended to have RSC values and FRiP scores within defined ranges (RSC between 1.5 and 2.5, FRiP ≥ 1%, **Figure 2 D and E**).

Although it is well established that no one single metric appears to reliably assess ChIP-seq quality (Landt et al. 2012), FRiP, which measures the fraction of reads falling within peak regions, is a useful and straightforward metric for evaluation of ChIP-seq success. In general, FRiP values correlate positively and linearly with the number of called regions, and most TF ChIP-seq datasets from ENCODE have a FRiP enrichment of 1% or greater when peaks are called using MACS with default parameters (Landt et al. 2012). In our study, the FRiP values aligned with our initial assessment of data quality such that all five of our datasets with reduced signal strength had FRiP scores ranging from 0-0.9%, thus suggesting a sub-par or failed immunoprecipitation. Conversely, all of our other study datasets had FRiP scores of ≥ 1%, consistent with successful immunoprecipitation. By contrast, other quality metrics we tested (normalized strand coefficient, Q-tag) did not seem to be able to distinguish between successful and unsuccessful immunoprecipitation (data not shown).

As a further test of our hypothesis that chromatin size is one determinant of ChIP-seq quality, we then examined whether these objective measures of quality were also related to chromatin size. For these ChIP-seq samples, the RSC score was found to be strongly negatively associated with chromatin size (**Figure 3A)**, and the samples at both the low and high extremes of the sizes were failures (F) or low passes (L).

**Figure 3:**
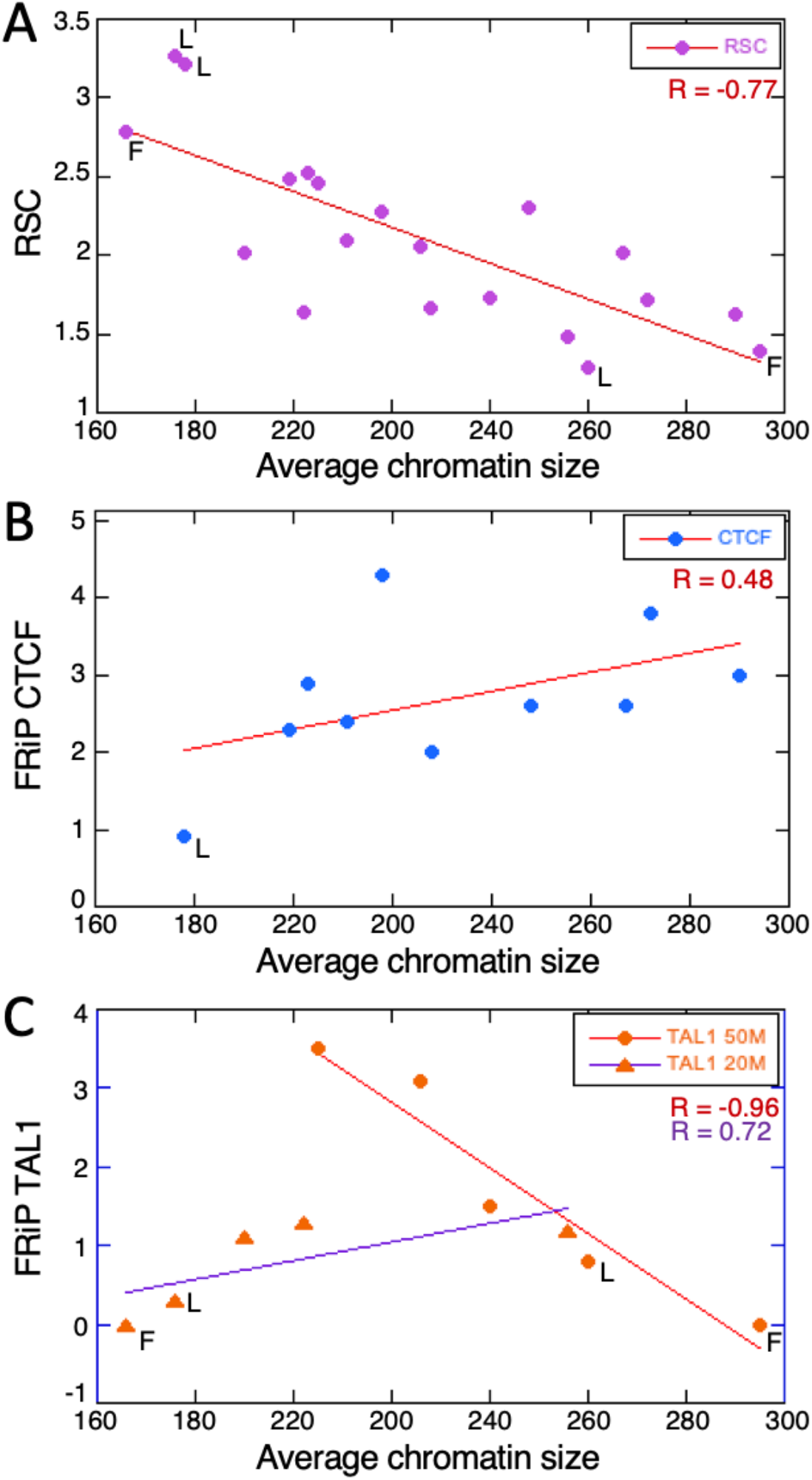
Relationship between chromatin size and objective quality metrics **A**. RSC vs chromatin size. **B**. FRiP (as percentage) vs average chromatin size for CTCF, **C**. FRiP percentage vs average chromatin size for TAL1. CTCF + TAL1 = purple, CTCF = blue, TAL1 = orange, 50M = circles, 20M = triangles, P = pass, L= low pass, F = fail.

The FRiP scores were also related to input chromatin size, with the scores for CTCF ChIP-seq datasets increasing with fragment length (**Figure 3B**). The scores for TAL1 ChIP-seq presented a more complex pattern with a consistent increase in the experimental group using 20M cells (R= 0.72), but a decline in FRiP scores at larger fragment sizes in the experimental group using 50M cells (R= -0.96, **Figure 3C**). Closer examination of the TAL1 datasets revealed that the datasets that had low FRiP scores also had the smallest (<180 bp) and largest (>250 bp) average chromatin length (**Figure 3C, Supplemental Table S1**).

A threshold of 1% for FRiP is frequently employed as a criterion for success of a ChIP-seq experiment (Landt et al. 2012), and that threshold fits well with our subjective visual assessments. As with the RSC assessment, the datasets at the extremes of the fragment size distribution tended to have FRiP scores indicative of poor quality. Overall, these analyses implicate the chromatin fragment size distribution (summarized as the mean) as an important determinant of success of a ChIP-seq experiment, and suggested the possibility that chromatin length may have some predictive value in determining ChIP-seq success.

### Retrospective analysis

To examine whether this relationship between average fragment size and success rate of ChIP-seq held for other samples, we did a retrospective analysis of ChIP-seq datasets that were performed in our laboratory. The inclusion criteria were as follows: 1) ChIP-seq experiments were performed in the same mouse erythroid cell line, G1E-ER4+E2, 2) the factor immunoprecipitated was either CTCF or TAL1, and 3) we had available unenriched input chromatin corresponding to the ChIP-seq library so that we could measure the average length of the chromatin used for the corresponding ChIP. A total of 12 retrospective datasets with independent chromatin sonications met these criteria. These ChIP-seq experiments were conducted over a variety of conditions, including different fixation methods, which confound our assessment of quality, but they provided an opportunity to determine whether a relationship between chromatin size and quality could still be detected. After assessing their quality by both by subjective inspection of ChIP-seq signals and by objective ENCODE quality metrics (**Supplemental Table S2**), we found that very low average chromatin size, a FRiP score below 1%, or a low RSC were associated with failure of the ChIP-seq experiment (**Figure 4**). Thus, this retrospective analysis further supports the trend that samples with small fragment sizes have poor results.

**Figure 4.**
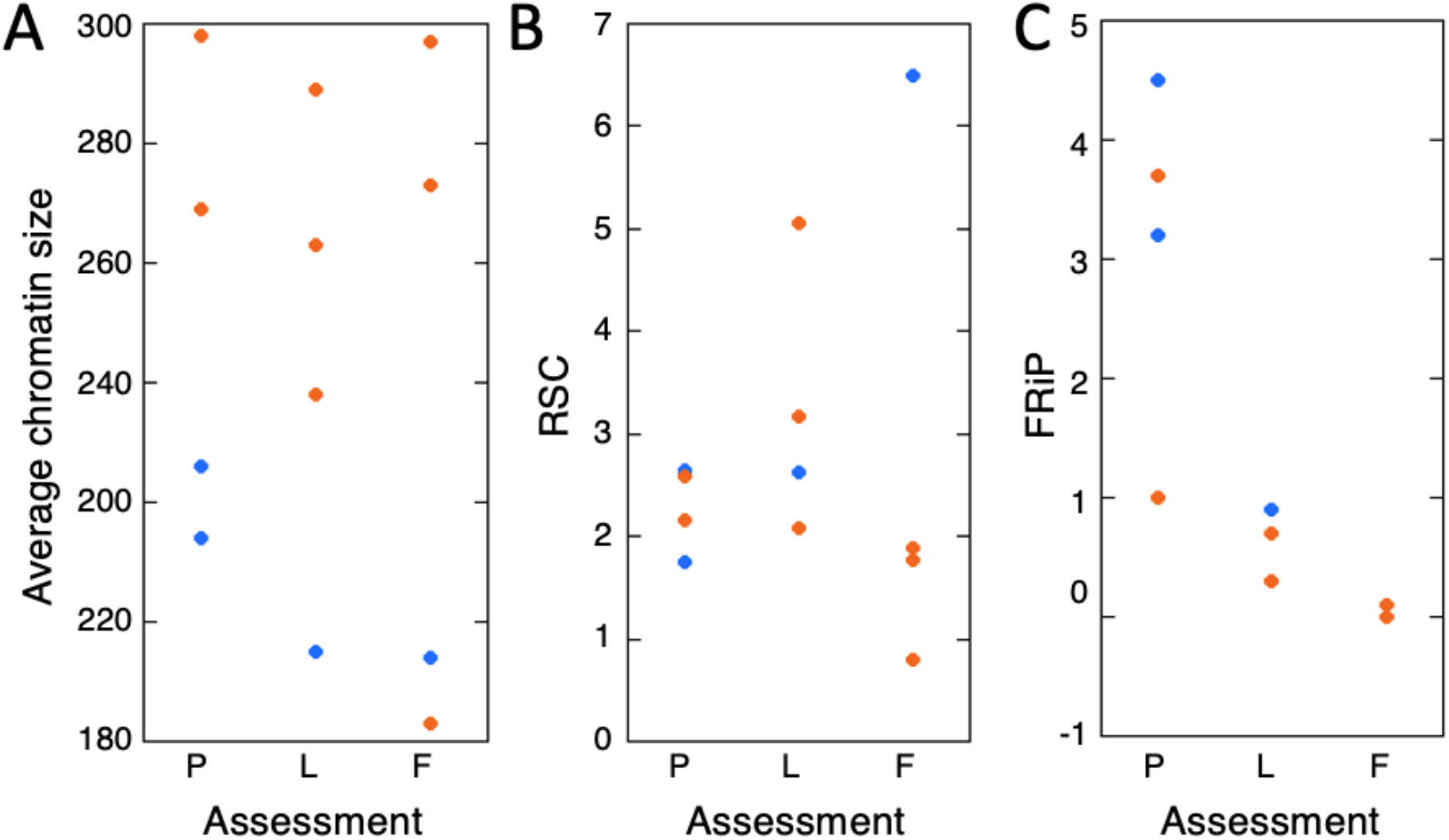
Retrospective datasets separated by subjective assessment. **A**. Average chromatin length. **B**. FRiP (as percentage) **C**. RSC. CTCF = blue, TAL1 = orange, 50M = circles, 20M = triangles, P = pass, L = low pass, F = fail.

To further test our hypothesis that ChIP-seq experimental success was dependent on the input chromatin size, we focused on the objective quality metric FRiP score. The study datasets and the retrospective datasets were combined (total of 32 experiments) and binned by the chromatin length (<200 bp, 200-250 bp, >250bp) for each TF. Our hypothesis predicts that the FRiP scores should be higher for experiments with chromatin sizes in specific ranges. The results support the hypothesis, as most samples of chromatin size <200bp have a low FRiP score indicative of failure, while experiments with samples of chromatin size between 200 and 250 bp have FRiP scores indicative of success (**Figure 5**). The differences between these two groups were significant for both CTCF (*p*<0.005, Student’s t-test) and TAL1 (*p*<0.05, Student’s t-test). Consistent with the previous analyses, almost all the TAL1 ChIP-seq experiments on samples with an average fragment size larger than 250 bp had mainly low FRiP scores, but CTCF ChIP-seq was successful in all metrics on samples in this larger size range. Despite this finding, the differences between >250 bp and 200-250 bp were not significant when examining all the data points for either CTCF or TAL1. Closer examination of the data revealed that one of the TAL1 ChIP-seq datasets with a mean size >250 bp had a very high FRIP score which appeared to skew the analysis. When the FRIP score for this unusual dataset was not included in the analysis, the differences between TAL1 200-250 bp and >250 bp were significant (p<0.05, Student’s t-test).

**Figure 5:**
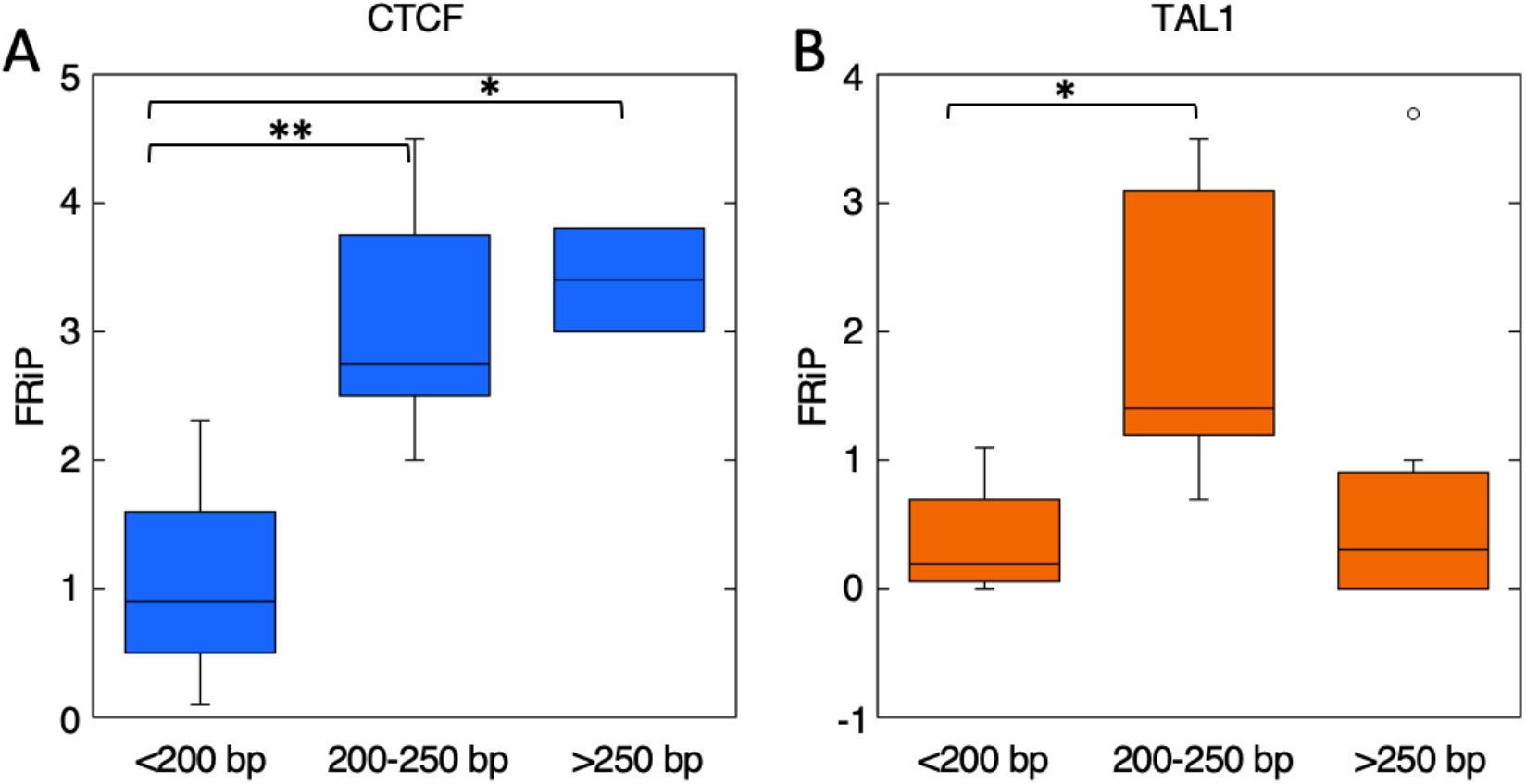
Relationship between chromatin size and FRiP score (as percentage) for study and retrospective datasets. **A**. CTCF. **B**. TAL1. CTCF = blue, TAL1 = orange, \**p*<0.05, \*\**p*<0.005.

Together, these results suggest that, under the conditions tested, sonication products between 200-250 bp yield the highest frequency of acceptable FRiP scores and successful ChIP-seq experiments. Further, while both CTCF and TAL1 are sensitive to over sonication, only TAL1 appears sensitive to under sonication. By contrast, CTCF appears to be more robust in ChIP-seq using larger MW chromatin (at least up to approximately 300 bp).

### Low input ChIP-seq

Obtaining reliable ChIP-seq results on TFs or other epigenetic features in cell types that can be purified only in small quantities would facilitate many studies, such as mechanisms regulating gene expression during cell differentiation. While some success has been reported for ChIP-seq in low cell input samples (Adli et al. 2010; Lara-Astiaso et al. 2014), these approaches have been primarily applied to modified histones, which are often present in greater abundance than TFs. We wanted to determine whether careful control of chromatin shearing would help facilitate low input ChIP-seq using standard methods. We performed ChIP-seq on 1M and 5M cells using an antibody against CTCF (**Figure 6A, Supplemental Table S3**), in contrast to our usual protocols involving 20M to 50M cells. We sheared groups of 1M cells for 3 or 4 cycles of 30 sec on, 30 sec off, and groups of 5M cells for 3, 5 or 7 cycles of 30 sec on, 30 sec off, assayed chromatin length for each sample, and performed ChIP-seq according to our standard protocol. We were unable to accurately determine the length of the chromatin samples from the 1M cell inputs using the Agilent Bioanalyzer because the DNA concentration was too low. The average sizes of the chromatin for the 5M cell samples were 329, 302, and 248 bp, respectively. As with previous samples, all datasets were first examined on the genome browser for the subjective assessment. As shown in **Figure 6A**, no signal was observed in any of the 1M cell samples or the 5M cell sample with an average chromatin length of 329 bp. By contrast, we observed signal tracks corresponding to CTCF occupancy in the 5M cell datasets with average chromatin lengths of 302 and 248 bp. Furthermore, the objective quantitative assessment of all the datasets revealed that only the 5M 302 bp and 5M 248 bp datasets had FRiP scores ≥1.0%.

**Figure 6:**
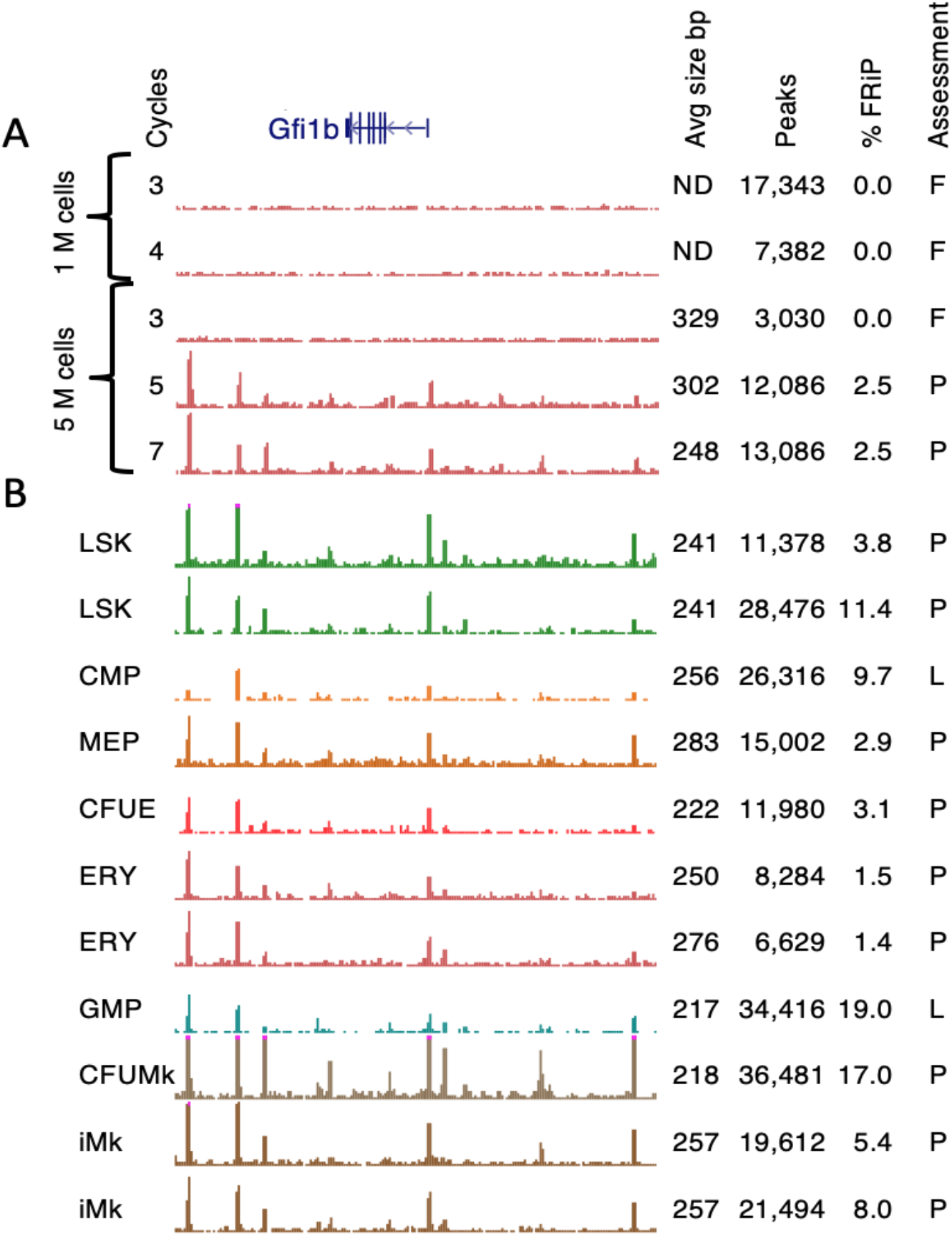
CTCF occupied sites at the *Gfi1b* locus in **A**. G1E-ER4+E2 cells, and **B**. Hematopoietic cell types. P = pass, L = low pass, F = fail, ND = not detectable.

Given that 5M cells were needed for successful conventional ChIP-seq for CTCF (if appropriate sonication conditions were employed), we felt that collecting a similar number of rare hematopoietic progenitor cells by sorting mouse bone marrow cells was feasible, despite the technical challenges. Thus, we leveraged our findings on low cell input to produce a set of CTCF ChIP-seq datasets in hematopoietic cell populations (**Figure 6B, Supplemental Table S3)**. These datasets include LSK (Lin-Sca1+Kit+, which includes hematopoietic stem cells or HSC), several multilineage progenitor cells (common myeloid progenitor cells or CMP, granulocyte monocyte progenitor cells or GMP, megakaryocyte erythrocyte progenitor cells or MEP), and committed cells of the major blood cell lineages at different stages of maturity (in the cases of erythroblasts or ERY and megakaryocytes or Mk). Due to progenitor cell rarity, it is challenging to obtain enough cells for a single ChIP-seq determination. For example, the isolation of 4.85M MEP cells required the bone marrow from 200 mice from a total of 10 flow cytometry sorts (**Supplemental Tables S3 and S4**). The progenitor cells were isolated and fixed in small batches (<1M cells, **Materials and Methods**) over the course of several months, and they were frozen at -80°C until enough cells were amassed to attempt a CTCF ChIP-seq experiment. Isolation procedures of this magnitude involve a significant time investment, and thus it is critical on the part of the experimentalist to do everything possible to increase the probability of ChIP-seq success.

As before, we used the Diagenode Biorupter 300 to shear fixed chromatin from the fixed hematopoietic cells, assayed the extent of shearing using the Agilent Bioanalzyer 2100, and then subjected the chromatin to immunoprecipitation and sequencing using antiserum against CTCF (**Figure 6B, Supplemental Table S3**). We first inspected each of the signal tracks in all 11 datasets in 8 different cell types across several loci, including the *Gfi1b* locus. We observed peaks in all 11 datasets across 8 different cell types examined, though there were notable differences in the signal to noise ratio between samples. Based on this initial subjective visual inspection, two of the 11 datasets were deemed as “Low pass” while the remaining datasets were classified as “Pass”. Examination of the objective metrics for these datasets revealed a large number of peaks and, in some cases, very high FRiP scores with all of the datasets having a FRiP score of at least ≥1%.

In summary, by rigorous attention to sonication conditions, we were able to obtain good quality CTCF binding profiles across a series of cells differentiating from multilineage progenitor cells to lineage-specific, maturing blood cells.

## Discussion

ChIP-seq has been used extensively across a broad spectrum of species, tissues, and cell types to interrogate the locations of TFs occupying specific sites in chromatin or mapping the profile of histone modifications in chromatin (Wold and Myers 2008, Rivera and Ren 2013). This powerful technique has moved studies of gene regulation to a global (genome-wide) scale, and the data produced by this method form the foundation for many efforts to develop coherent, integrated models for gene regulation (Ching et al. 2018; Zhou et al. 2019; Xiang et al. 2020). However, the technique is not uniformly successful for all samples, and even when apparently successful, the resulting ChIP-seq datasets vary widely in quality (Marinov et al. 2014; Devailly et al. 2015). A major factor impacting success of the ChIP-seq is the antibody directed against the chromatin-associated factor (TF or histone modification). Effective antibodies not only must be highly specific, but they also must recognize epitopes that may not be sufficiently exposed in the fixed chromatin. These requirements are difficult to evaluate prior to performing the ChIP-seq experiment, since many antibodies that are effective for other applications, such as Western blots, are not effective in ChIP-seq (Egelhofer et al. 2011). Thus, the only assay for effectiveness of an antibody is to actually do the ChIP-seq experiment, which requires expenditure of resources and time, and of course it is frustrating when the experiment fails. Such failures are frequent for commercially available antibodies (Egelhofer et al. 2011; Savic et al. 2015). Sophisticated tagging approaches have been developed to enable the use of antibodies known to be effective in a context where the level of antigen is close to the natural level (Savic et al. 2015). These approaches help to generate more uniform results for TFs expressed in cell lines or organisms amenable to the targeted genetic engineering required for tagging. Other factors also contribute to the success of a ChIP-seq experiment, some of which are directly under the control of the experimentalist. One illustration of high technical variation arises when a particular lot of a commercial antibody preparation is used successfully in ChIP-seq for one biosample, but then a subsequent experiment fails, even though the same lot of antibody was used in another preparation of that same biosample. The studies in this paper were developed to better understand the technical contributors to success of a ChIP-seq experiment, beyond the well-known issues with antibody quality.

We discovered that the size distribution of the sonicated chromatin was an important factor in the ChIP-seq procedure, with higher successes observed for chromatin with an average size in a range of approximately 200 to 250 bp. We conclude that monitoring the level of chromatin sonication is one way to improve ChIP-seq quality and reproducibility. Indeed, we would argue that in situations with failure and success for the same antibody in the same biosample, it would be wise to re-evaluate the chromatin length for these samples. In such instances, it is *critical* to test the actual chromatin to be used in a ChIP-seq experiment.

The best sonication conditions should be determined empirically for each cell type, number of cells, and ideally any treatments applied to those cells that could cause changes in cell morphology. While some parameters for sonication may be expected to be predictable, the many factors impacting the final chromatin size distribution mean that an empirical approach is needed. For example, as expected, we observed that samples with larger numbers of cells required more sonication cycles than those with fewer cells to achieve a given size range. However, it is should be noted that very low numbers of cells may actually require more sonication cycles than larger numbers of cells because the incidence of a sound wave interacting with the chromatin decreases as the overall chromatin size decreases, relative to the wavelength of the insonifying sound. Thus, it may require additional time to have sufficient interaction between the sound wave and the chromatin (Espana et al. 2014). While not specifically addressed in the current presentation, it is reasonable to expect the amount of required sonication to vary by tissue and cell type. To facilitate these empirical determinations, we provide a protocol for determining chromatin size distribution in the desired range in **Supplemental Figure S1 and S2**.

We demonstrated an effect of chromatin size distribution on success of ChIP-seq experiments for two TF targets. While the experiments for both tended to fail at small chromatin sizes, the effects of larger chromatin sizes differed, with the CTCF ChIP-seq experiments being less sensitive to large sizes of chromatin, compared to TAL1 ChIP-seq. One hypothesis to explain the differences in effects of larger chromatin sizes is a difference in the exposure of the antigens. Specifically, one can conjecture that TAL1 antigens are more sequestered in longer chromatin fragments, whereas the CTCF antigens may be more exposed in those longer fragments. Indeed, one may expect the details of the relationship between chromatin fragment size and ChIP-seq success to vary among different TF targets, but working within a range with frequent success (200-250 bp for average fragment sizes) is a good starting point for experiments.

In agreement with previous work (Landt et al. 2012), we found that FRiP scores of at least 1% to be strongly associated with successful ChIP-seq experiments targeting TFs. We also noted some experimental results with very high FRiP scores, e.g. 17% to 19%. Such very high scores do not necessarily indicate an exquisite dataset. For instance, in a dataset with a low signal to noise ratio, a peak-calling algorithm may call an excessive number of false positive “peaks”. Those overcalled peaks are included when counting the number of reads assigned to peaks, and thus, the FRiP score can be artificially inflated.

In addition to improving the success rate for conventional ChIP-seq experiments, careful control of chromatin shearing may help facilitate ChIP-seq experiments on low numbers of input cells. We showed that, with rigorous attention to sonication conditions, we were able to obtain good quality CTCF binding profiles across a series of cells differentiating from multilineage progenitor cells to lineage-specific, maturing blood cells. This series includes rare progenitor cells that have been difficult to interrogate for binding profiles of specific TFs. These maps of binding by CTCF across progenitor and lineage-specific cells should be useful for multiple studies of the roles of this important architectural protein during blood cell differentiation.

## Materials and Methods

### Cell culture methods

G1E-ER4 cells were grown in IMDM media + 15% FBS, kit ligand, and erythropoietin in a standard tissue culture incubator at 37°C with 5% CO^2^ as described (Weiss et al. 1997). To induce erythroid maturation, G1E-ER4 cells were treated with 10^−8^ mol/L β-estradiol for 24 hours (G1E-ER4+E2). Cells were harvested by centrifugation at 500 x *g* for 5 min at 4°C and washed once in 1xPBS.

### Isolation of hematopoietic progenitor cells

All primary hematopoietic cell populations were enriched from 5-8 week old C57BL6 male mice. LSK, CMP, MEP, GMP, CFU-E, ERY, CFU-MK, and iMK populations were harvested and isolated from bone marrow (BM) using the following cell isolation markers as described (Heuston et al 2018, **Supplemental Table S4**). LSK: CD4-, CD8-, IL7Ra-, CD11b-, Ly6g-, CD45R-, Ter119-, cKit+, Sca1+ CMP: CD4-, CD8-, IL7Ra-, CD11b-, Ly6g-, CD45R-, Ter119-, cKit+, Sca1-, CD34+, CD16/32-GMP: CD4-, CD8-, IL7Ra-, CD11b-, Ly6g-, CD45R-, Ter119-, cKit+, Sca1-, CD34+, CD16/32+ MEP: CD4-, CD8-, IL7Ra-, CD11b-, Ly6g-, CD45R-, Ter119-, cKit+, Sca1-, CD34-, CD16/32-CFUE: CD4-, CD8-, IL7Ra-, CD11b-, Ly6g-, CD45R-, Ter119+, CD44+, FSCHigh ERY: CD4-, CD8-, IL7Ra-, CD11b-, Ly6g-, CD45R-, Ter119+, CD44+, FSCLow CFUMk: CD4-, CD8-, IL7Ra-, CD11b-, Ly6g-, CD45R-, Ter119-, Sca1-, CD41+, CD61+, cKit+ iMK: CD4-, CD8-, IL7Ra-, CD11b-, Ly6g-, CD45R-, Ter119-, Sca1-, CD41+, CD61+, cKit-

### Chromatin Immunoprecipitation (ChIP)

For CTCF or TAL1 in G1E-ER4+E2, either 1M, 5M, 20M, or 50M cells in 1x PBS were crosslinked for 10 min by adding formaldehyde at a final concentration of 0.4% and glycine was added at a final concentration of 125 mM to quench cross-linking. Cells were washed in 1xPBS and stored at -80°C until needed.

For CTCF in LSK, CMP, MEP, GMP, CFUE, ERY, CFUMk, iMk, cells were fixed in 0.4% formaldehyde (16% methanol-free, Thermo Scientific) for 15 minutes before quenching in 125 mM glycine for 5 minutes. Cells were washed in 2X PIC (Roche mini-tabs, 1 tab in 5 ml = 2X) and stored at -80°C until needed.

Cells were then lysed (10 mM Tris-HCl, pH 8.0, 10 nM NaCl, 0.2% NP40) for 10 min on ice, washed once in 1x PBS, followed by nuclear lysis (50 mM Tris-HCl 8.0, 10 mM EDTA, 1% SDS) for 10 min on ice. Chromatin was then diluted further with Immunoprecipitation Buffer (20 mM Tris-HCl, pH 8.0, 2 mM EDTA, 150 mM NaCl, 1% Triton X-100, 0.01% SDS) and a 1x Protease Inhibitor Cocktail set V, EDTA-free (Calibiochem, La Jolla, CA). A Diagenode Bioruptor 300 was used to shear samples in cycles of 30 sec on, 30 sec off sonication at high output power at 4°C for the desired number of cycles. For optimization of sonication, see the protocol detailed in **Supplemental Figure S1**.

Sonicated chromatin was pre-cleared overnight at 4°C with 8 μg of normal goat IgG (Santa Cruz Biotechnology, Santa Cruz, CA; sc2028) for TAL1 or 8 μg of normal rabbit IgG (sc2027) for CTCF. Separately, 5 μg of TAL1 antibody (Santa Cruz Biotechnology, sc12984x, lot B2511) or 5 μL of CTCF antiserum (MilliporeSigma, 07-729) were pre-bound to protein G agarose beads overnight at 4°C. For binding, pre-cleared chromatin was added to the antibody:bead complex and incubated with rotation at 4°C for 4 hours. 200 μL of pre-cleared chromatin was saved for use as input. After binding, the beads were washed with Wash Buffer I (20 mM Tris-HCl, pH 8.0, 2 mM EDTA, 50 mM NaCl, 1% Triton X-100, 0.1% SDS), High Salt Wash Buffer (20 mM Tris-HCl, pH 8.0, 2 mM EDTA, 500 mM NaCl, 1% Triton X-100, 0.1% SDS), Wash Buffer II (10 mM Tris-HCl, pH 8.0, 1 mM EDTA, 250 mM LiCl, 1% NP40, 1% deoxycholate), and 1x TE. DNA:protein complexes were then eluted from beads with Elution Buffer (1% SDS, 100 mM NaHCO3). Reverse crosslinking of immunoprecipitated chromatin was accomplished by the addition of NaCl to ChIP and input samples, followed by incubation overnight at 65 °C with 1μg RNase A. To remove proteins, each sample was treated with 6 μg Proteinase K for 2 hours at 45 °C. Finally, immunoprecipitated DNA was then purified using the Qiagen PCR Purification Kit (Qiagen, Germantown, MD).

### Illumina Library Preparation for ChIP-Seq

All samples, including input, were processed for library construction for Illumina sequencing using Illumina’s TruSeq ChIP-seq Sample Preparation Kit according to manufacturer’s instructions. The DNA libraries were sequenced on the HiSeq 2000 or NextSeq 500 as indicated (**Supplemental Tables S1, S2, S3, S5)** using Illumina’s kits and reagents as appropriate.

### ChIP-seq data processing

The sequencing reads were mapped to mouse genome assembly mm10 using Bowtie (0.12.8), and then filtered to remove chrM, unplaced chromosomes, and unmapped reads. The alignment is converted to bam format using Samtools 0.1.8. MACS (1.3.7.1) is used to generate the wiggle tracks and call peaks. UCSC’s wigToBigWig program is used to convert the wiggle file to a bigWig.

## Data Access

Tables with ChIP-seq samples, statistics on numbers of reads and mapped reads, and quality metrics are in the **Supplemental Tables S1, S2, S3, S5**. Data are deposited in the NCBI Gene Expression Omnibus (GEO; https://www.ncbi.nlm.nih.gov/geo/), GEO accession number [_ in progress_]

## Acknowledgements

We thank D.E.C. for insight into scattering of sound from submerged objects. This work was supported by grants from the National Institutes of Health NIDDK R24 DK106766 and NHGRI U54 HG006998.

## References

Adli M, Zhu J, Bernstein BE. 2010.Genome-wide chromatin maps derived from limited numbers of hematopoietic progenitors. Nat Methods 7: 615–618.

Browne JA, Harris A, Leir SH. 2014.An optimized protocol for isolating primary epithelial cell chromatin for ChIP. PLoS One 9: e100099.

Ching T, Himmelstein DS, Beaulieu-Jones BK, Kalinin AA, Do BT, Way GP, Ferrero E, Agapow PM, Zietz M, Hoffman MM et al. 2018. Opportunities and obstacles for deep learning in biology and medicine. J R Soc Interface 15.

Davis CA, Hitz BC, Sloan CA, Chan ET, Davidson JM, Gabdank I, Hilton JA, Jain K, Baymuradov UK, Narayanan AK et al. 2018. The Encyclopedia of DNA elements (ENCODE): data portal update. Nucleic Acids Res 46: D794–d801.

Devailly G, Mantsoki A, Michoel T, Joshi A. 2015.Variable reproducibility in genome-scale public data: A case study using ENCODE ChIP sequencing resource. FEBS Lett 589: 3866–3870.

Diagenode. 2012.The Ultimate Guide for Chromatin Shearing Optimization with Bioruptor® Standard and Plus. Diagenode User’s Guide.

Dogan N, Wu W, Morrissey CS, Chen KB, Stonestrom A, Long M, Keller CA, Cheng Y, Jain D, Visel A et al. 2015. Occupancy by key transcription factors is a more accurate predictor of enhancer activity than histone modifications or chromatin accessibility. Epigenetics Chromatin 8: 16.

Egelhofer TA, Minoda A, Klugman S, Lee K, Kolasinska-Zwierz P, Alekseyenko AA, Cheung MS, Day DS, Gadel S, Gorchakov AA et al. 2011. An assessment of histone-modification antibody quality. Nat Struct Mol Biol 18: 91–93.

Espana AL, Williams KL, Plotnick DS, Marston PL. 2014.Acoustic scattering from a water-filled cylindrical shell: Measurements, modeling, and interpretation. The Journal of the Acoustical Society of America 136: 109–121.

Khoja H, Smejkal G, Krowczynska A, Herlihy JD. 2019.Optimizing Sample Fixation and Chromatin Shearing forImproved Sensitivity and Reproducibility of ChromatinImmunoprecipitation. Covaris Application Note.

Landt SG, Marinov GK, Kundaje A, Kheradpour P, Pauli F, Batzoglou S, Bernstein BE, Bickel P, Brown JB, Cayting P et al. 2012. ChIP-seq guidelines and practices of the ENCODE and modENCODE consortia. Genome Res 22: 1813–1831.

Lara-Astiaso D, Weiner A, Lorenzo-Vivas E, Zaretsky I, Jaitin DA, David E, Keren-Shaul H, Mildner A, Winter D, Jung S et al. 2014. Immunogenetics. Chromatin state dynamics during blood formation. Science 345: 943–949.

Marinov GK, Kundaje A, Park PJ, Wold BJ. 2014.Large-scale quality analysis of published ChIP-seq data. G3 (Bethesda) 4: 209–223.

Savic D, Partridge EC, Newberry KM, Smith SB, Meadows SK, Roberts BS, Mackiewicz M, Mendenhall EM, Myers RM. 2015.CETCh-seq: CRISPR epitope tagging ChIP-seq of DNA-binding proteins. Genome Res 25: 1581–1589.

Weiss MJ, Yu C, Orkin SH. 1997.Erythroid-cell-specific properties of transcription factor GATA-1 revealed by phenotypic rescue of a gene-targeted cell line. Mol Cell Biol 17: 1642–1651.

Wilson NK, Foster SD, Wang X, Knezevic K, Schütte J, Kaimakis P, Chilarska PM, Kinston S, Ouwehand WH, Dzierzak E et al. 2010. Combinatorial transcriptional control in blood stem/progenitor cells: genome-wide analysis of ten major transcriptional regulators. Cell Stem Cell 7: 532–544.

Wilson NK, Schoenfelder S, Hannah R, Sánchez Castillo M, Schütte J, Ladopoulos V, Mitchelmore J, Goode DK, Calero-Nieto FJ, Moignard V et al. 2016. Integrated genome-scale analysis of the transcriptional regulatory landscape in a blood stem/progenitor cell model. Blood 127: e12–23.

Xiang G, Keller CA, Heuston E, Giardine BM, An L, Wixom AQ, Miller A, Cockburn A, Sauria MEG, Weaver K et al. 2020. An integrative view of the regulatory and transcriptional landscapes in mouse hematopoiesis. Genome Res 30: 472–484.

Zhou J, Schor IE, Yao V, Theesfeld CL, Marco-Ferreres R, Tadych A, Furlong EEM, Troyanskaya OG. 2019.Accurate genome-wide predictions of spatio-temporal gene expression during embryonic development. PLoS Genet 15: e1008382.

