## Supplemental Figures for "Effects of sheared chromatin length on ChIP-seq quality and sensitivity"

### Supplemental Figure S1

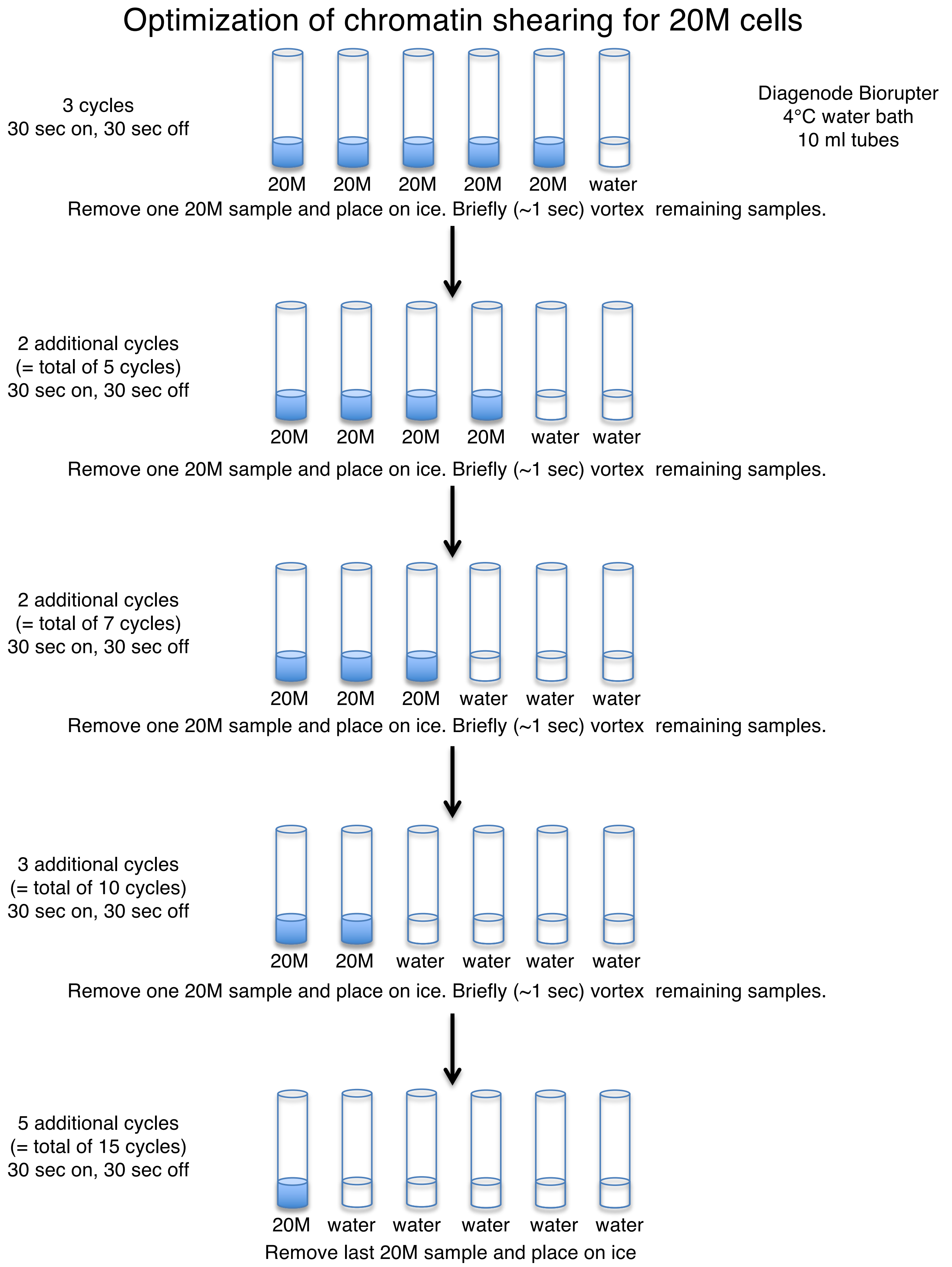

#### Supplemental Figure S1 continued

### Optimization of chromatin shearing for 20M cells

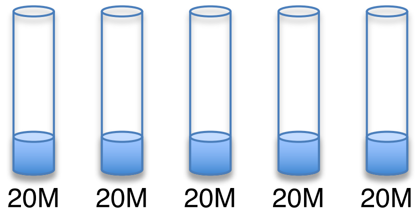


Sheared chromatin on ice

Either preclear chromatin o/n
or
Freeze at -20°C for short term storage
or
Freeze at -80°C for long term storage

Remove 50 µL of each condition for testing*

Bring volume up to 200 µL with ChIP Elution Buffer
Add: 16 µl of 5M NaCl
1 µl of 1 mg/ml of RNase A
Incubate o/n at 65°C

<u>ChIP Elution Buffer</u>
100 mM $NaHCO_3$
1% SDS

3µL of 20 mg/mL proteinase K to each tube and incubate for 2 hours at 45°C.

Purify samples using a Qiagen PCR purification kit
Elute samples in 100 µl of <u>Qiagen</u> EB solution according to kit protocol.

<u>Qiagen EB solution</u>
(from Qiagen kit)

Run samples on Agilent Bioanalyzer.
Region 100-500 bp and determine average length.
Proceed with ChIP-seq and determine which size range may be best for your factor of interest.

Once you have determined an acceptable range, you do not need to repeat titration, but you should always check your samples to make sure they are in the predetermined acceptable range.

*You can use less than 50 µl, however, we always increase the volume to 200 µl with ChIP elution buffer and process samples in the same way that we process ChIP samples for reverse cross-linking. If you sample less than 50 µl, please also remember to decrease your elution volume from Qiagen column accordingly.

### Supplemental Figure S2

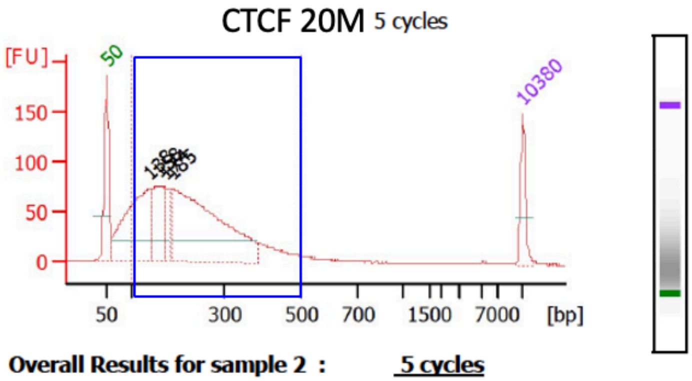


**Peak table for sample 2 :** **5 cycles**

| Peak | | Size [bp] | Conc. [ng/µl] | Molarity [nmol/l] |
|---|---|---|---|---|
| 1 | ◀ | 50 | 8.30 | 251.5 |
| 2 | | 138 | 21.05 | 231.2 |
| 3 | | 158 | 9.77 | 93.9 |
| 4 | | 174 | 3.32 | 28.9 |
| 5 | | 185 | 37.36 | 305.8 |
| 6 | ▶ | 10,380 | 4.20 | 0.6 |

**Overall Results for sample 2 :** **5 cycles**

Number of peaks found: 4

Area 1: 921.4

**Region table for sample 2 :** **5 cycles**

| From [bp] | To [bp] | Area | % of Total | Average Size [bp] | Size distribution in CV [%] | Conc. [ng/µl] | Co lor |
|---|---|---|---|---|---|---|---|
| 100 | 500 | 921.4 | 84 | 228 | 39.8 | 64.74 | |

**Supplemental Figure S2:** Representative Aglient Bioanalzyer 2100 tracing of a chromatin sample using an Agilent DNA 7500 kit. Following sonication, the average size of each chromatin sample was determined by measuring the average length within a 100-500 bp window.
